# Collective dynamics of DNA methylation during ageing

**DOI:** 10.1101/2024.12.15.628564

**Authors:** Aida Hashtroud, Marc Bonder, Oliver Stegle, Ferdinand von Meyen, Wolf Reik, Steffen Rulands

## Abstract

Ageing is the decline of physiological function over time. Statistical models termed ageing clocks can predict chronological and biological age from the longitudinal time evolution of DNA methylation. Here, we show that DNA methylation ageing is also manifest in how pairs of genomic loci evolve with respect to each other with age. Using sequencing data from a range of tissues in mouse we show that genomic correlations in DNA methylation during ageing are characterised by an enrichment of correlations between sites that are roughly 500 bp apart. We trace the origin of this behaviour to collective dynamics in the boundaries of CpG islands. We derive a simple, biophysical model that explains the origin due to a tilt in the competition between methylating and demethylating processes and an ensuing wetting-like phenomenon. Using Chip-seq data, we argue for a molecular mechanism based on a PRC2-dependent dilution of H3K27me3 during ageing. Our work gives a new perspective on epigenetic ageing, highlighting the importance of correlative as opposed to longitudinal dynamics.

## 1. Introduction

Ageing refers to the gradual decline of physiological functions in an organism over time [1]. This involves multiple processes at various biological scales, such as cellular senescence, stem cell depletion, telomere shortening, and mitochondrial dysfunction [2]. Epigenetic changes, including DNA methylation, histone modifications, and chromatin remodelling, are also closely linked to ageing [3, 4, 5]. Among these, DNA methylation, which primarily occurs at cytosines in a CpG context, is one of the key epigenetic mechanisms associated with ageing (Fig. 1A). During early development, de-novo DNA methylation is established by DNMT3A/B enzymes, while DNA replication generates hemimethylated sites that are corrected by a complex involving DNMT1 and UHRF1[6, 7, 8, 9]. In the absence of DNMT1 or UHRF1, DNA methylation can be gradually diluted across multiple cell divisions [10, 11]. DNA methylation marks can also be actively removed by TET1/2/3 enzymes and through the base excision repair pathway [6, 7].

**Figure 1:**
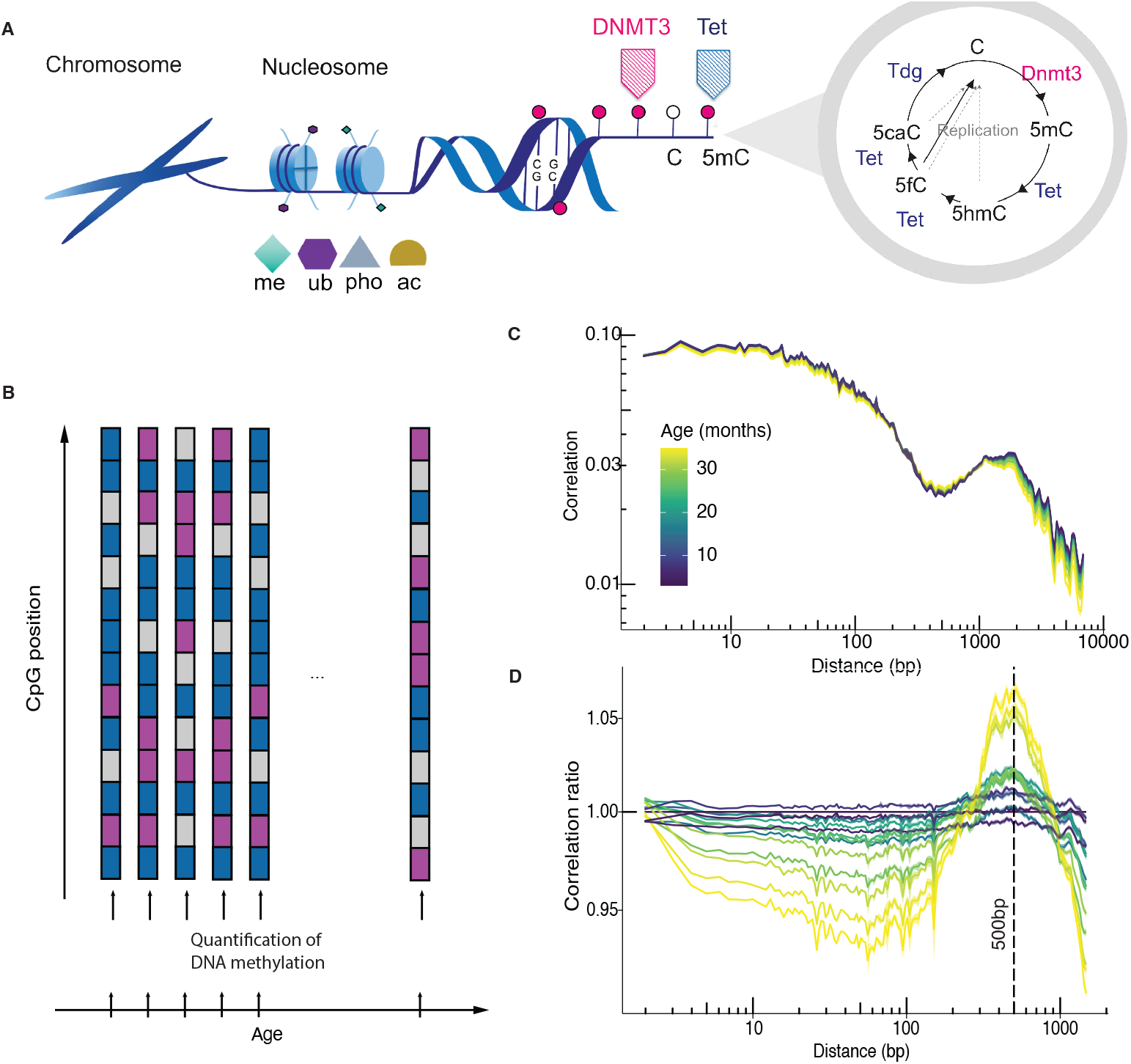
**A** Schematic of basic epigenetic modifications of the DNA and chromatin. **B** Schematic of a longitudinal quantification of DNA methylation during ageing using whole-genome DNA methylation sequencing. **C** Correlation functions of genome-wide DNA methylation states for different ages in mouse blood (data taken from [16]). Shaded regions are standard errors. Distances were grouped into XXX bins and the average correlation in each bin is shown. **D** Correlation ratio with respect to the youngest age. Shaded areas are 95% confidence intervals obtained with bootstrapping. Colours as in **C**.

Both, DNA methylation and histone modifications exhibit systematic changes with age [12, 13, 14]. Statistical models based on the linear combination of DNA methylation states at numerous CpG sites can predict chronological and biological age across different tissues and species [15, 16, 17, 18]. These models, called DNA methylation ageing clocks, are trained on DNA methylation measurements at individual CpGs using bulk sequencing methods, which pool information across many cells [5]. Recent studies have also observed systematic DNA methylation changes at the single-cell level[19, 20]. It remains unclear which regulatory process, if any, is captured by DNA methylation ageing clocks [21]. Recent works suggest that DNA methylation ageing clocks measure the accumulation of stochastic variations over time[22, 23, 24, 25]. Consistent with this idea, DNA methylation levels tend to converge toward intermediate levels with age [16]. CpG islands, typically low in methylation, gradually gain methylation [26], while highly methylated regions associated with heterochromatin, repetitive elements, and transposons tend to lose methylation over time [2, 27].

DNA methylation ageing clocks capture the longitudinal evolution of a set of individual CpGs over time. Here, we take an orthogonal perspective and show that the progression of epigenetic ageing is also reflected in how DNA methylation states co-evolve in relation to each other over time. Specifically, drawing on bisulfite sequencing data from various tissues, we show that epigenetic ageing is characterized by an increase in correlations between CpG sites that have a distance of roughly 500 bp. By deriving a biophysical model we show that this observation originates from collective dynamics between antagonistic enzyme molecules in the shores of CpG islands. This dynamics gives rise to a wetting-like phenomenon of enzymes involved in the demethylation of CpG islands. Based on the analysis of Chip-seq experiments in combination with bisulfite sequencing experiments we argue for a molecular mechanism, in which an ageing-associated dilution of PRC2 activity leads to a tilt in the competition between DNMT3A and TET enzymes in the shores of bivalent CpG islands.

## 2. Results

### 2.1. Collective changes in DNA methylation during ageing

To understand collective changes in DNA methylation during ageing, we analyzed a range of bisulfite sequencing experiments on various tissues in mouse during ageing. These include blood, liver, cortex, heart, lung, and kidney [16, 18]. To quantify the co-evolution of DNA methylation states at different genomic positions, we calculated the so-called two-point correlation functions of DNA methylation states. Mathematically, the correlation function is defined as *C*(*l*) = ⟨*m*_*i*_*m*_*i*+*l*_⟩ − ⟨*m*_*i*_⟩⟨*m*_*i*+*l*_⟩, where *m*_*i*_ and *m*_*i*+*l*_ are the DNA methylation values at genomic positions *i* and *i* + *l*, respectively. Intuitively, the correlation function quantifies the degree to which the DNA methylation states at sites with a genomic distance *l* are linked. Figure 1C shows correlation functions over different ages for blood. While there is a systematic change with age, the correlation function also depends on features that are independent of age. These stem, for example, from the sequence-dependence of DNA methylation levels.

Earlier work has shown that uncoordinated noise is an important factor in predicting the time evolution of DNA methylation during ageing along with a general drift toward inter-mediary DNA methylation levels [10, 25]. Because this entropy-driven change is expected from the laws of thermodynamics we here focus on systematic changes in the correlations between DNA methylation states. To this end, we noted that while the correlation function *C*(*l*) is affected by uncoordinated noise, the ratio of correlation functions taken at different ages, *r*(*l*) = *C*(*l*)*/C*^′^(*l*), only captures changes that are systematically correlated in the genome. Uncoordinated changes in DNA methylation levels lead to a distance-independent correlation ratio, *r*(*l*) = const., while changes that are coordinated in the genome lead to a distance-dependent correlation ratio. Using this approach, we found that correlations in DNA methylation between genomic positions decrease, on average, for distances smaller than roughly 200 bp and larger than 1000 bp (Fig. 1D, Supplementary Fig. S1A). Significantly, for a characteristic distance of 500 bp, the DNA methylation state of CpGs becomes increasingly correlated during ageing.

To investigate the mechanistic origin of why sites with this particular distance become increasingly correlated during ageing, we first asked whether this behaviour was prevalent across the genome or whether it was limited to specific genomic regions. We reasoned that the average size of CpG islands in the mouse genome is 1045± 5 bp (SEM). The distance of 500 bp could therefore be related to the typical distance of a CpG in CpG islands to their boundaries, called the CpG island shores. To test this, we computed the correlation ratio separately for CpG islands and CpG poor regions (Fig. 2A). For CpG-poor regions the correlation ratio is independent of the distance, which is in agreement with uncoordinated noise. By contrast, CpG islands exhibit an increase in DNA methylation values of sites with a distance of 500 bp. We obtained analogous results when using the Pearson correlation (Supplemental Fig. S1B). We then sorted CpG islands by size and found that the characteristic distance at which DNA methylation correlations increase during ageing scales with the size of CpG islands (Fig. 2B), which further supports a central role of collective DNA methylation dynamics in CpG island shores.

**Figure 2:**
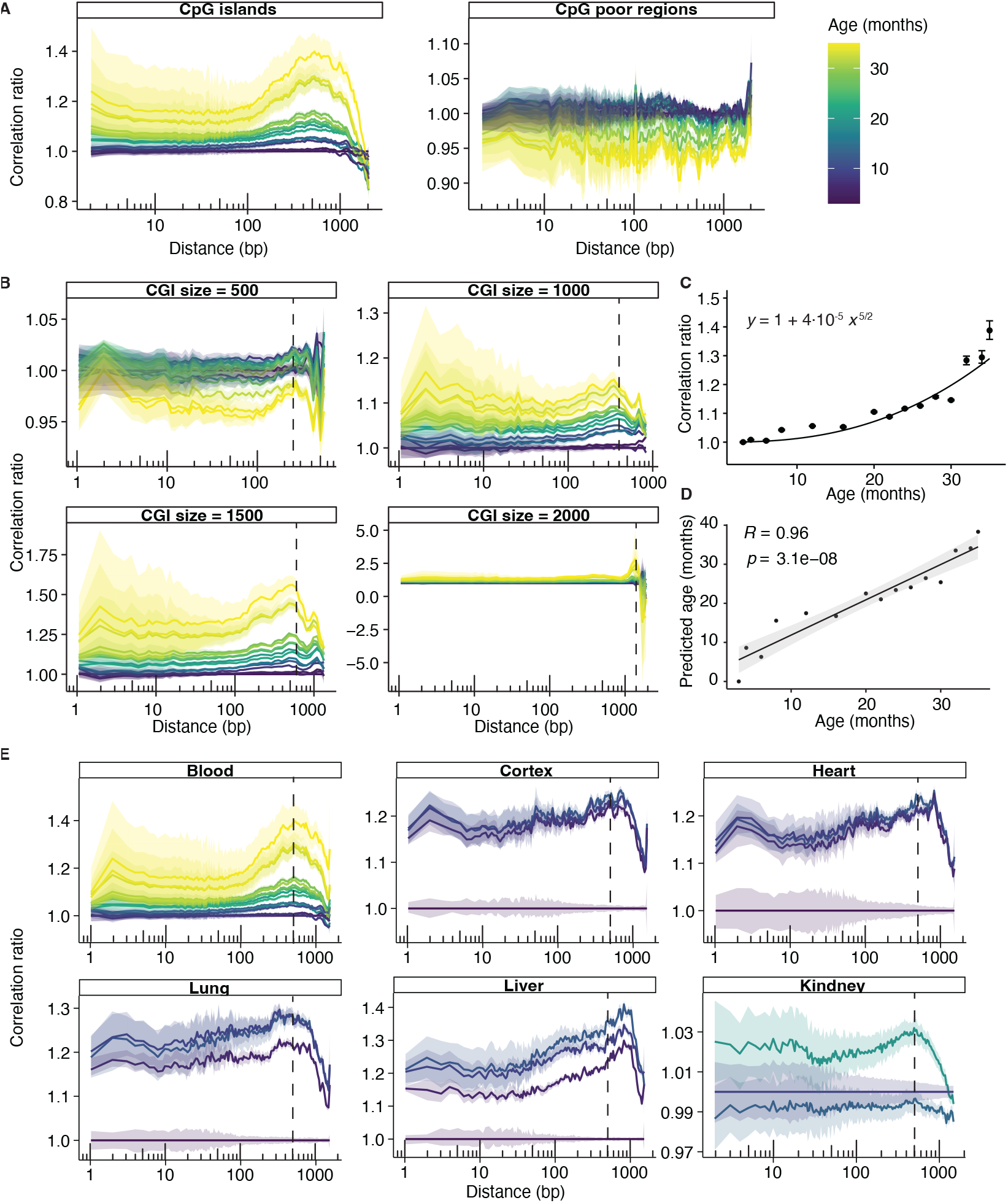
**A** Correlation ratio for CpG islands (left) and the remainder of the genome (right). Shaded areas in this and all other figures denote 95% confidence intervals obtained from bootstrapping. Colour indicates age and is consistently used throughout all panels in 7this figure. **B** Correlation ratio for CpG islands sorted by size. The position of the maximal enrichment of correlation scales (dashed vertical lines) with the CpG island size. **C** Average correlation ratio over distances between 450 bp and 550 bp. The solid line depicts a heuristic model as indicated in the figure. **D** Prediction of chronological age using the model from **C.** The shaded area denotes 95% confidence intervals of the linear regression line (solid line). **E** Correlation ratios for different tissues using data from Refs. [16], [18], and [28]. Vertical dashed lines indicate 500 bp.

The systematic change in DNA methylation correlations during ageing can, in principle, be used to construct DNA methylation ageing clocks. Constructing such an ageing clock is not the aim of the present work but to demonstrate the feasibility thereof we defined a statistical model that describes the overall trend in the gain in correlations at distances between 450 and 550 bp (Fig. 2C). This model is a rough heuristic description of the overall trend and we did not employ any parameter fitting. Such a model can then be inverted and used to predict chronological age with good accuracy (Fig. 2D).

When filtering for CpG islands we found an analogous coordinated change in DNA methylation in all tissues we analysed, which include Blood, Cortex, Heart, Lung, Liver, and Kidney (Fig. 2C) as well as intestinal stem cells (Supplementary Fig. S1C). To a lesser degree, we observed the same behaviour in a more noisy data set of FACS-sorted T cells, but not in B cells. Taken together, these analyses suggest that DNA methylation dynamics on the shores of CpG islands underlie the collective behaviour quantified in Fig. 1D. These changes in the shores of CpG islands are systematic and could be used to predict chronological age (Fig. 2C).

### 2.2. A minimal biophysical model

The observation in Fig. 2A is a reflection of a systematic, collective process and cannot be explained by the entropic accumulation of noise alone [25]. To demonstrate this, we initialized stochastic simulations with DNA methylation values taken at the youngest age present in Ref. [16]. We then evolved DNA methylation states over time following purely stochastic dynamics and quantified correlation functions and correlation ratios in CpG islands as in Fig. 2A. Figures S2A,B show that, as expected from the definition of the correlation ratio, stochasticity alone cannot explain the observations in Fig. 2A. To understand how such dynamics can come about we used biophysical modelling. With this model, we aimed to show how the collective behaviour depicted in Figs. 2A and 2C can arise as the result of the self-organization of biophysical processes in CpG island shores. To this end, we proceeded in two steps: First, rather than quantifying all individual enzymatic processes happening in CpG islands, we aimed to derive the simplest possible model that can predict the data. Then, in a second step, we derived a molecular mechanism that explains the parameters constituting this model in detail.

We began by considering the dynamics of enzymes that mediate DNA methylation and demethylation in CpG islands. CpG islands are largely lowly methylated. Demethylation is mediated by TET enzymes that bind to the centre of the CpG island [29, 7, 6, 30]. DNMT3A enzymes are de-novo methyl transferases that localize in the genomic regions outside of the CpG island [31, 31, 30]. To describe the kinetics of these enzymes in the CpG island shores, we described the DNA in and around a CpG island as a one-dimensional lattice, where each site can be occupied by a TET or DNMT3A enzyme [32]. Both types of enzymes have been shown to catalyze their own binding [33, 34, 35]. The shores of CpG islands are regions of competition between these antagonistic enzymes as has been shown by perturbation experiments [30, 36]. To further constrain the model, we also applied the condition of detailed balance that relates the binding and unbinding rates of enzymes in thermodynamic equilibrium. With this, the model is defined by the binding energies of TET enzymes in the presence of other TET enzymes, *J*^*T T*^, DNMT3A enzymes in the presence of other DNMT3A enzymes, *J*^*DD*^, and the binding energy of one enzyme in the presence of the other, *J*^*DT*^.

One-dimensional models tend to be strongly sensitive to the presence of fluctuations. To model these fluctuations correctly, we therefore needed to consider the effect of the three-dimensional structure of the genome on the one-dimensional enzyme kinetics [32]. To this end, we let the binding affinities be distance-dependent in a way that is given by the internal end-to-end distance probability of the DNA [37]. This probability decays with the distance between two CpG sites to the power of an exponent *γ*. Theoretical studies predict exponents that range between 1*/*3 for space-filling compacted chromatin and 2 for a self-avoiding random walk [32]. Hi-C experiments indicate distance-dependent values smaller than 1 for the short distances relevant for CpG islands [38]. Because these exponents describe long-range genomic interactions, the feedback of the three-dimensional chromatin structure leads to an averaging out of fluctuations and allows stable genomic domains to form (Methods).

With this framework, we hypothesized that ageing is associated with a tilt in the competition between DNMT3A and TET enzymes. Mathematically, this is described by a function that monotonically increases during ageing and subtracts energy *h* from each DNMT3A-bound site. We will discuss the molecular mechanism behind this shift in competition further below.

The changes that occur during ageing are much slower than the time it takes for the enzyme binding and unbinding processes to reach a steady state. We may therefore consider the tilt in the competition between TET and DNMT3A to be quasistatic, such that at each time point during aging the molecular processes have equilibrated. This allowed us, in the next step, to derive a thermodynamic description of the model (Methods). To this end, we next systematically coarse-grained the lattice model. This process involved defining adjacent groups of CpG sites which we iteratively increased in size and then averaged over molecular fluctuations [39]. As a result of this procedure, we obtained a thermodynamic free energy of the form

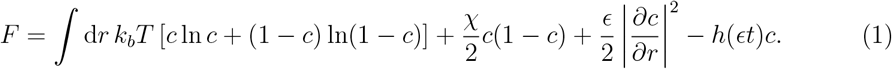

Here, *c* is a rescaled concentration of bound DNMT3A enzymes, and *χ* = *J*^*T T*^ +*J*^*DD*^−2*J*^*DT*^ is a parameter that quantifies the degree to which both enzymes tend to bind in the vicinity of enzymes of the same kind. Further, *k*_*B*_ is the Boltzmann constant and *T* is the temperature. *ϵ* denotes the rate with which ageing occurs and which we assume to be much slower than any individual molecular processes in the CpG island, *ϵ* ≪ 1. At a given age, the most likely enzyme binding profile then simply follows from minimizing the free energy.

The benefit of a thermodynamic description is that it allowed us to interpret the model in biophysical terms. In this free energy, the first term describes the mixing entropy of DNMT3A and TET enzymes, the second term their competitive binding, the third term an effective diffusive spreading, and the fourth term the effect of ageing. Free energy densities of the form (1) are known to describe spinodal decomposition or phase separation phenomena in a wide variety of settings, such as in subcellular condensates [40]. Phase separation is usually impossible in one-dimensional systems but here facilitated by an interplay between processes in the 1d and the 3d genome. In the context of DNA methylation, this means that DNMT3A and TET proteins spatially separate along the genome via a self-organization process.

### 2.3. Model predictions

The expression of the model in terms of a free energy also provides a convenient way to explore a wide range of model behaviours. First, the model predicts several phase transitions as a function of the parameters *χ* and *h*, which describe the strength of enzyme competition and the tilt in favour of TET, respectively (Fig. 3C). For low values of both of these parameters, the model predicts a mixing of the binding positions of DNMT3A and TET enzymes. The model also allows for a spinodal phase, in which DNMT3A and TET domains separate into spatially distinct genomic regions. This happens if both enzymes preferentially bind in the vicinity of enzymes of the same kind and *χ* therefore exceeds a threshold value. In between the mixed and spinodal phase is a binodal regime, in which the separation of both domains is long-lived but will ultimately vanish. Although it is not possible to quantify the biological values of these parameters directly, the experimental observation of competition between DNMT3A and TET islands in CpG island shores [31, 36] suggests that biologically plausible parameters are in the spinodal phase.

**Figure 3:**
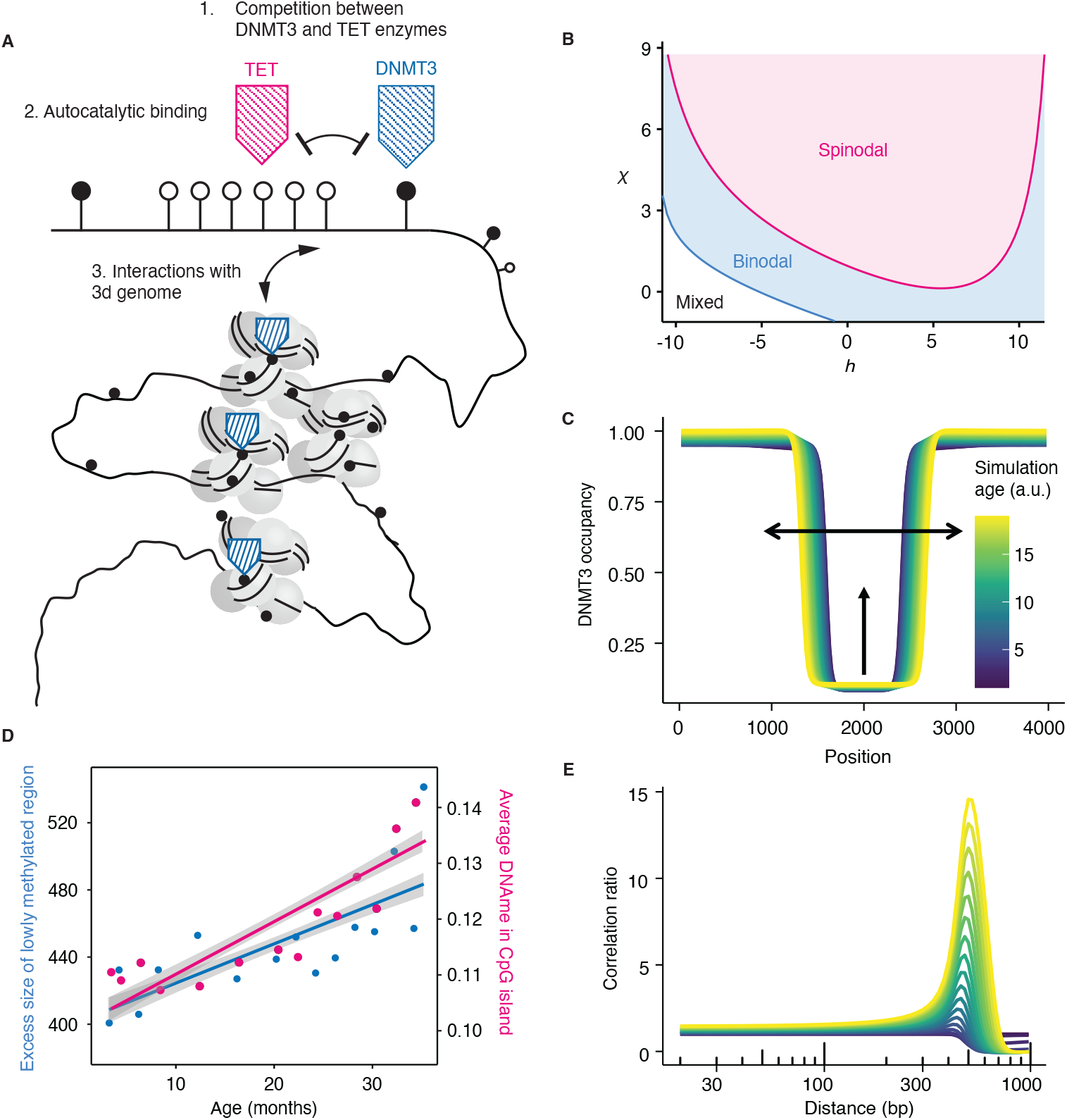
**A** Schematic depicting the processes constituting the model. **B** Phase diagram obtained from minimizing the free energy, Eq. 1. The blue line separates the mixed from the binodal regime and the red line the binodal and the spinodal (phase separating) regime. **C** Simulation of DNMT3 binding profiles for different ages. Shown are quasi-steady states obtained from minimizing the free energy at each age. Ageing was implemented as a linearly increasing function with *τ*. Arrows indicate the spreading of the DNMT3A-low region and the gain of DNMT3 binding in the CpG island. **D** Size of the lowly methylated region in CpG islands (blue) and average DNA methylation in CpG islands (pink) at different ages. Lines are regression lines and shaded areas indicated 95% confidence intervals. **E** Correlation ratios obtained from the same simulations as in C.

The free energy in Eq. (1) also allows for predicting the time evolution of enzyme binding profiles during ageing (Methods). To this end, we initialized the model Eq. (1) such that DNMT3A occupies regions far away from the centre of the CpG island positioned around *r* = 0. In the spinodal phase, the lowly methylated region in the centre of the CpG island with high TET and low DNMT3A occupancy spreads out with a velocity that is determined by the age-dependence of the change in competition, *h*. Simultaneously, TET occupancy in this region decreases with an implied gain of DNA methylation in the CpG island. This behaviour and the underlying free energy resemble partial wetting phenomena of liquids on surfaces [41].

We next tested whether we could observe this behaviour in experimental data on DNA methylation during ageing. To this end, we analyzed rrBS-seq experiments on blood ageing [16] and quantified at each time point both the average DNA methylation level in CpG islands and the size of the associated hypomethylated region (Fig. 3D). In agreement with previous studies [26, 42, 43, 23] we found that DNA methylation levels in CpG islands tend to increase over time. Simultaneously, we found that the size of the hypomethylated region increased with age.

We then asked whether the model was able to predict the correlation function in Figs. 2A and 2D. To this end, we ran stochastic simulations of the model incorporating both noise associated with the enzyme kinetics and variability in the genomic sizes of CpG islands but did not perform any kind of parameter tuning apart from setting the size of the CpG island to 1000 bp on average (Methods). These simulations indeed show the emergence of enriched correlations at a distance of 500 bp as observed in the experiment (Fig. 4E).

**Figure 4:**
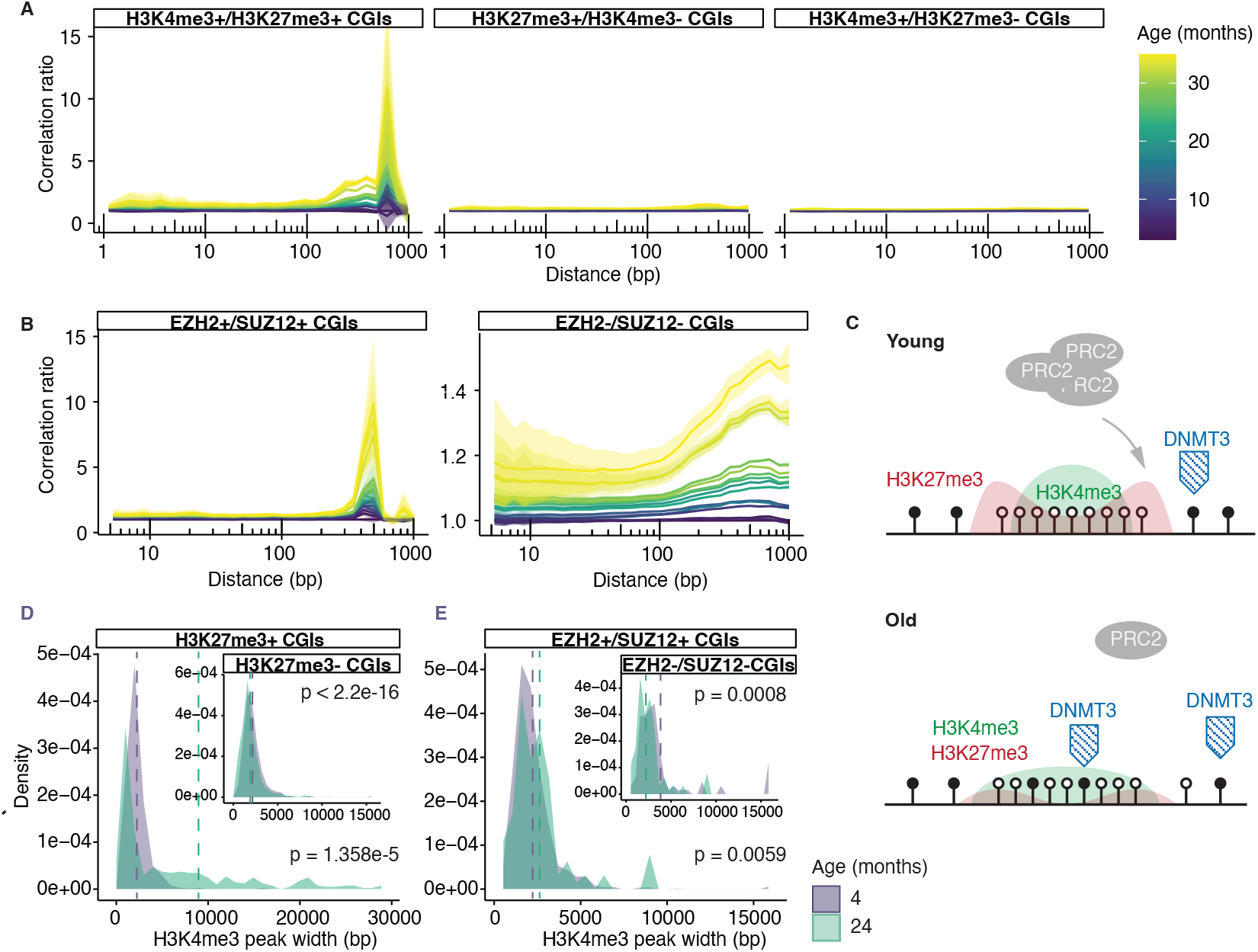
**A** Correlation ratio for CpG islands depending on overlap with H3K4m34 or H3K27me3 annotations obtained from Chip-seq experiments. **B** Correlation ratios for CpG islands depending on the overlap with binding regions of the EZH2 and SUZ12 components of the PRC2 complex. **C** Cartoon depicting the molecular mechanism underlying the model. **D** Probability density of the widths of H3K4me3 peaks overlapping with H3K27me3 peaks for young and old mice. Inlay: probability density of the widths of H3K4me3 peaks not overlapping with H3K27me3 peaks. **E** Probability density of the widths of H3K4me3 peaks overlapping with SUZ12 and EZH2 peaks for young and old mice. Inlay: probability density of the widths of H3K4me3 peaks not overlapping with SUZ12 and EZH2 peaks. P-values in D and E were obtained using Wilcoxon tests.

Taken together, our minimal mechanistic model can predict key features of the collective evolution of DNA methylation during ageing: the simultaneous expansion of lowly methylated regions with the increase of DNA methylation in the same region. It further predicts the changes in the co-evolution of DNA methylation in pairs of CpG sites as encoded in the correlation function.

### 2.4. Molecular mechanism

While this model gives a mechanistic explanation for the observed collective changes in DNA methylation during ageing, it cannot inform about the detailed molecular processes that comprise the tilt in the competition between DNMT3A and TET enzymes. We therefore used further profiling data for the different molecular players interacting with DNMT3A and TET enzymes in CpG islands. While these analyses further clarify the molecular origins of the parameters in Eq. (1), a mathematical model incorporating these molecular details would not yield distinguishable predictions on DNA methylation correlations.

As a first hypothesis for the molecular mechanism underlying the tilt in enzyme competition during ageing, we reasoned that DNMT3A has been shown to mutate in blood during ageing [44]. It has also been found that DNMT3A mutant cells are clonally advantageous, such that their proportion in blood samples increases over time [45]. Similar observations have been made for TET enzymes [30]. A changing proportion of clones with altered DNMT3A or TET binding rates could potentially explain the observed changes in DNA methylation correlations. However, while these might be potential explanations for a tilt in an effective DNMT3A-TET competition in blood they do not explain the same observations in other tissues not subject to clonal hematopoiesis. We, therefore, sought for a mechanism that is not tissue-dependent.

To this end, we first asked whether we could get a more specific identification of the CpG islands that show the collective DNA methylation changes in Figs. 2A and 2D. Specifically, we asked whether these changes were more or less pronounced in the presence of different histone tail modifications that interact with DNA methylation. The active histone mark H3K4me3 is associated with CpG islands and correlated with gene activity for CpG islands overlapping with promoter regions [46]. It is deposited and maintained during replication by the H3K4 methyltransferase COMPASS complex [47, 48] which again represses DNA methylation [49]. The repressive histone mark H3K27me3 is associated with inactive genes silenced by the polycomb repressive complex 2 (PRC2) [50], which again recognizes TET1 [51]. Both H3K4me3 and H3K27me3 suppress the binding of DNMT3A [30]. H3K4me3 and H3K27me3 not only have opposite effects on gene activity, but they are also mutually repressive in that the catalytic activity of PRC2, which mediates H3K27me3, is inhibited by H3K4me3 [52, 53, 54, 55]

Using annotations from Chip-seq experiments on mouse blood [12] we found that the enrichment in DNA methylation correlations was specific to CpG islands that were enriched both for H3K4me3 and H3K27me3, so-called bivalent CpG islands (Fig. 4A). This pattern in the data was much less expressed for CpG islands annotated with either H3K4me3 or H3K27me3, but not both. In bivalent CpG islands, H3K4m3 tends to be located in the centre and H3K27me3 closer to the shores [56, 57].

Bivalent CpG islands have been identified to be targets of PRC2 [58]. The role of PRC2 in epigenetic ageing has been well documented: PRC2 has been found to be diluted during ageing [59]. PRC2 to is also known to interact with DNA methylation, in that PRC2 binding regions tend to become hypermethylated during ageing [60, 42, 43, 23, 61]. This is despite a biochemical antagonism between PRC2 and DNA methylation [62, 63, 64, 65, 66], .To test the role of PRC2 in the collective DNA methylation dynamics we calculated DNA methylation correlations separately for bivalent CpG islands that are PRC2 targets and CpG islands that are not. To this end, we used Chip-seq data for two of the components of the PRC2 complex, SUZ12 and EZH2 [67]. Indeed, we found that the enrichment of DNA methylation correlations was specific to binding regions of both SUZ12 and EZH2 (Fig. 4B), but for SUZ12 individually (Fig. S3A,B).

Taken together, this suggests a molecular mechanism for the hypothesized tilt in the competition between DNMT3A and TET enzymes and ultimately the systematic, correlative changes in DNA methylation during ageing (Fig. 4C): the reduction of PRC2 activity leads to a dilution of H3K27me3, potentially due to cell replication [68]. Indeed it has been reported that H3K27me3 is reduced during aging [12]. This has two consequences: First, an expansion and simultaneous dilution of H3K4me3 and, consequently, an expansion of the lowly methylated region with a simultaneous gain of DNA methylation [66]. This is in line with work that showed that a dilution of PRC2 increases DNMT3A occupancy in the CpG islands [51]. In support of this mechanism, we analyzed H3K4me3 Chip-seq data from mouse blood [12]. We calculated the H3K4me3 peak width for old (24 months) and young (4 months old) mice separately for peaks overlapping with several other annotations. We found that the average width of H3K4me4 peaks overlapping with H3K27me3 increased four-fold between young and old mice (Fig. 4D). By contrast, H3K4me3 peaks that did not overlap with H3K27me3 peaks did not increase in width but slightly decreased (Fig. 4D, inlay). Similarly, H3K4me3 peaks overlapping with SUZ12 and EZH2 peaks increased on average in width during ageing (Fig. 4E), while H3K4me3 peaks not overlapping with those annotations did not increase in width during ageing but slightly decreased (Fig. 4E, inlay). Taken together, this shows that, as predicted, the width of H3K4me3 regions increases during ageing and that this spreading is associated with the simultaneous presence of both H3K4me3 and H3K27me3 and binding of the PRC2 complex. Therefore, the change in the competition between DNMT3A and TET enzymes could be a reflection of a parallel change in the competition between the COMPASS and PRC2 complexes [69, 49], associated downstream effects on H3K4me3 and H3K27me3 dynamics, DNMT3A and TET binding and ultimately DNA methylation.

## 3. Discussion

DNA methylation is highly predictive of chronological age. DNA methylation ageing clocks quantify the longitudinal evolution of a large number of CpG sites. Here, we showed that ageing is not only associated with the longitudinal time evolution of the DNA methylation state of CpG sites but also with changes in how CpG sites are correlated across extended genomic distances. These correlative changes are manifest in a characteristic increase of correlations of the DNA methylation states of CpGs at a distance of around 500bps, which are associated with the shores of PRC2-targeted CpG islands.

Our results explain several seemingly unrelated findings in epigenetic ageing research: First, the observation of correlated ageing dynamics in CpG island shores may explain why CpGs located in CpG island shores are enriched in the set of sites picked up by some DNA methylation ageing clocks [18]. The tilt in the competition between DNMT3A and TET enzymes in CpG island shores may also explain why DNMT3 mutant cell lines are assigned older chronological ages compared to wildtype cells [70].

Just as the longitudinal evolution of CpG sites over time also their genomic correlations can be used to build accurate ageing clocks. Here, we only provided proof of principle and instead focused on developing a biophysical model that mechanistically explains the collective dynamics of DNA methylation during ageing. The biophysical model captures essential ingredients of the collective DNA methylation dynamics during ageing, such as the competition between antagonistic enzymes in CpG island shores, and can also make quantitative predictions that are tested in sequencing data. The biophysical model necessarily does not capture the full complexity of the molecular processes. By using genomic annotations for the binding of a number of proteins and histone modifications involved in the regulation of bivalent CpG islands we resolve the molecular processes underlying the parameters of the biophysical model. Importantly, a model that would incorporate all of these processes would make indistinguishable predictions on DNA methylation correlations but would be statistically less strong due to its higher complexity.

The relation between PRC2 and DNA methylation changes in CpG islands, and, in particular, the counterintuitive gain of DNA methylation upon a decrease in PRC2 activity, has been subject to debate [66]. In our model, the gain in DNA methylation in CpG islands upon a decrease in PRC2 activity arises naturally as a result of molecular self-organization processes. Biologically, these ultimately trace back to stability H3K4me3 upon replication as opposed to H3K27me3. Physically, the model suggests a notable analogy to the physics of wetting processes on surfaces. In this mathematical analogy, enzymes located in the centre of CpG islands spread out after the dilution of repressive factors closer to the shores. This spreading is predicted to quantitatively resemble the behaviour of a liquid exhibiting surface tension. Liquid-like behaviour of protein-DNA complexes has been observed before in the phenomenon of protein-DNA co-condensation [71]. In this view, the simultaneous spreading of the lowly methylated domains and the gain of DNA methylation in that domain have a unified explanation as a result of a self-organization process of different molecular players in CpG islands. Whether proteins associated with PRC2-targeted CpG islands indeed have liquid-like properties is an interesting question for future research.

## Acknowledgements

We thank F. J. Meigel, C. Modes, M. Huch, F. Jülicher for their helpful feedback and all members of the involved groups for critical discussions. This work was supported by BBSRC (BBS/E/B/000C0421, BBS/E/B/000C0422, Core Capability Grant to W.R.), a Wellcome Trust Investigator Award (210754/Z/18/Z to W.R.), a King’s Prize Fellowship (King’s Health Partners, Wellcome Trust and London Law Trust to F.v.M.), ETH Zurich core funding (F.v.M.), and the European Research Council (ERC) under the European Union’s Horizon 2020 research and innovation programme (Starting Grant 803491 to F.v.M.; Starting Grant 950349 to S.R.).

## 4. Methods

### 4.1. Analysis of rrBS-seq data

Data was processed in the same way as in the original publications. Correlation functions were calculated using custom C code and further processing was performed in R4.0.2. The connected correlation function is defined as *C*(*l*) = ⟨*m*_*i*_*m*_*i*+*l*_⟩™⟨*m*_*i*_⟩⟨*m*_*i*+*l*_⟩, where *m*_*i*_ and *m*_*i*+*l*_ are the DNA methylation values at genomic positions *i* and *i* + *l*, respectively. Averages were performed over all genomic positions *i*, chromosomes, and replicates. To calculate correlation functions over genomic annotations, positions *i* were first filtered for overlaps with a given annotation before averages were taken.

To determine the size of the lowly methylated region linked to each CpG island, we used custom R code in which we first computed rolling averages of DNA methylation values with a window size of 10 CpGs. We then defined lowly methylated genomic segments as consecutive CpGs having a value of the rolling average less than 50% of the average DNA methylation level in the CpG island and the CpG island shores, i.e. regions of size 5000 bp adjacent to the CpG islands. The size of the lowly methylated region was then defined as the difference between the end of the last and the start of the first lowly methylated segment in a genomic region including the CpG island and its shores. Fig. 3D shows the difference between this value and the age-independent size of the genomic region annotated as a CpG island.

### 4.2. Theory

#### 4.2.1. Microscopic model

To define a simple biophysical model describing the time evolution of DNA methylation correlations during ageing, we begin by considering a one-dimensional lattice model comprising binary state variables. Each site in the lattice corresponds to a CpG site, which is either occupied by DNMT3 or a TET enzyme. We assign indices to the lattice sites using the notation *i* = 1, …, *N*, and the overall configuration of the lattice is represented by the set of all variables, *σ* = {*s*_1_, …, *s*_*i*_, …*s*_*N*_}. In this notation, the variable *s*_*i*_ = 1 signifies that site *i* is occupied by DNMT3A enzyme while *s*_*i*_ = 0 signifies TET occupation. The effect

of the geometry of the three-dimensional genome was taken into account for by making all energies dependent on the distance |*i* − *j*| with a specific distance dependence of the form 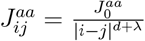 [32]. For biological reasons but also to ensure the existence of the thermodynamic limit these interactions should further be cutoff exponentially as |*i* − *j*| → ∞ [72]. Here, 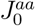 is a constant coefficient that sets the strength of the interaction. The value of *d* is 1.

The evolution of a given initial configuration of this system is described by a master equation of the general form,

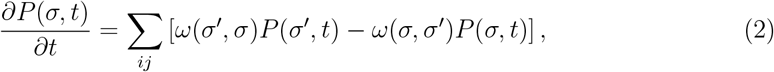

where *ω*(*σ, σ*^′^) is the transition rate from configuration *σ* to *σ*^′^. To reduce the number of free parameters we assume that the system is, at any given age, in thermal equilibrium. This implies that the molecular relaxation time to equilibrium is much faster than the systematic changes to the model parameters during ageing. At any given age quasi-equilibrium state at each age is given by the Boltzmann factor, *ω*(*σ, σ*^′^) ∝ *e*^−*βδH*^; where *β* = 1*/*(*k*_*B*_*T*) and *H* is the Hamiltonian. The Hamiltonian then reads

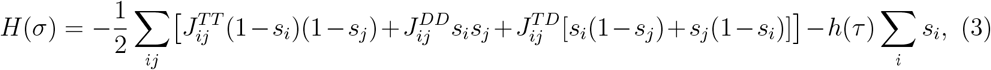

with *τ* = *ϵt*. Note that the first summation in the Hamiltonian includes all sites in the lattice to account for long-range interaction. The first summation can also be understood as exchange dynamics describing the interactions between DNMT3A and TET enzymes, while the second sum describes a reduction in the binding affinity of DNMT3A.

We can identically rewrite the Hamiltonian in Eq. (3) in terms of the interaction energy between site *i* and the rest of the system, in configuration *σ*, if site *i* is occupied by the 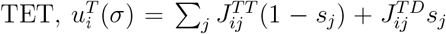, and analogously for 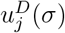. The transition rates then read

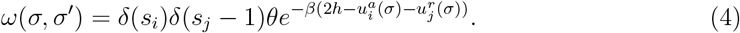

where *θ* is a characteristic exchange rate.

#### 4.2.2. Coarse-graining of the microscopic model

To obtain a description in terms of a free energy we follow the steps in Ref. [39]. To this end, we use a coarse-graining procedure in which we define a coarse-grained configuration 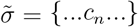. For the distance regome, where interactions follow a power law with an exponent smaller than 1 mean-field theory is correct. To also incorporate longer distances we defined coarse-grained concentration *c*_*n*_ is the average concentration of DNMT3 in compartment *n*,

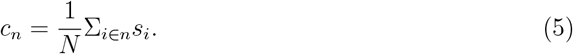

and *N* is the number of sites in each compartment. With this, we may rewrite the master equation in terms of a Fokker-Planck equation for the coarse-grained compartments,

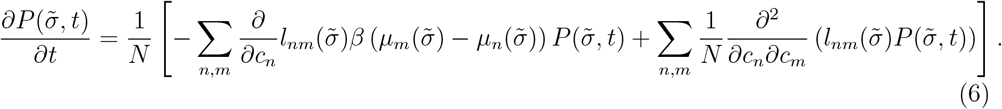

where *µ*_*n*_ is different in chemical potentials between a DNMT3 bound and a TET bound state in compartment *n*. Further, 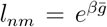 with 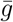 being the total chemical potential in compartments *n* and *m*. We can then express the dynamics in compartment *n* in terms of a Langevin equation of the form

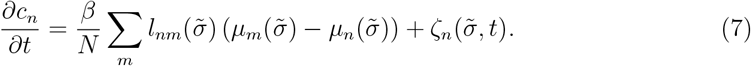

*ζ*_*n*_ is a multiplicative Gaussian noise as a function of the coarse-grained configuration, with the following moments,

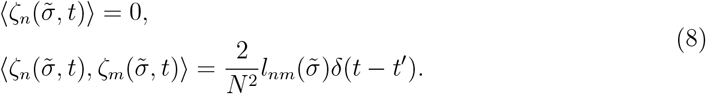

Equation (7) is equivalent to the conventional Cahn-Hilliard equation [73, 74] with noise which takes the form

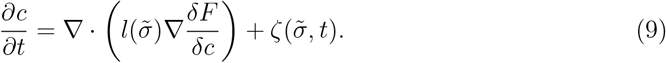

Note that we absorbed the dependence on *N* in the definition of the concentrations. The connection to the Cahn-Hilliard equation can be made explicit by deriving the total free energy,

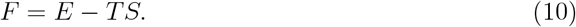

The configurational entropy of the system is [75],

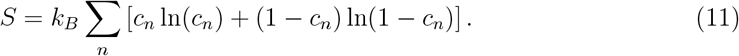

With this, the total free energy reads

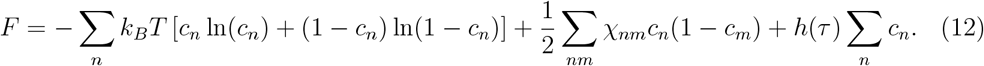

where we defined

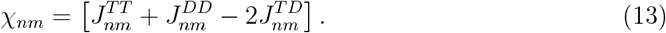

The continuum form of the free energy (12) is obtained by noting that

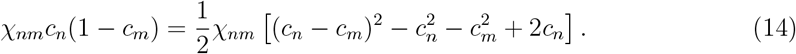

In the continuum limit, the total free energy then reads,

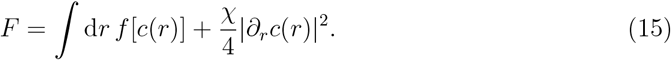

where *χ* = _*m*_ *χ*_*nm*_, and the nonlinear local part of the free energy per unit volume, *f*, is,

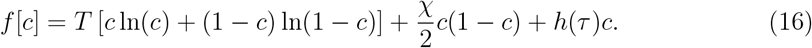

### 4.2.3. Computation of the phase diagram and the time evolution of the binding profile

To compute the phase diagram in Fig. 3 we numerically minimized the free energy for combinations of the parameters *χ* and *h* using custom code in Matlab. To compute the time evolution of DNMT3 binding profiles we obtained quasi-static solutions of Eq. (7) for a given value of *h*. Shown in Fig. 3C are quasi-static solutions for increasing values of *h* between 1 and 40 which were obtained numerically using an Euler scheme in Matlab.

## Supplementary figures

**Figure S1:**
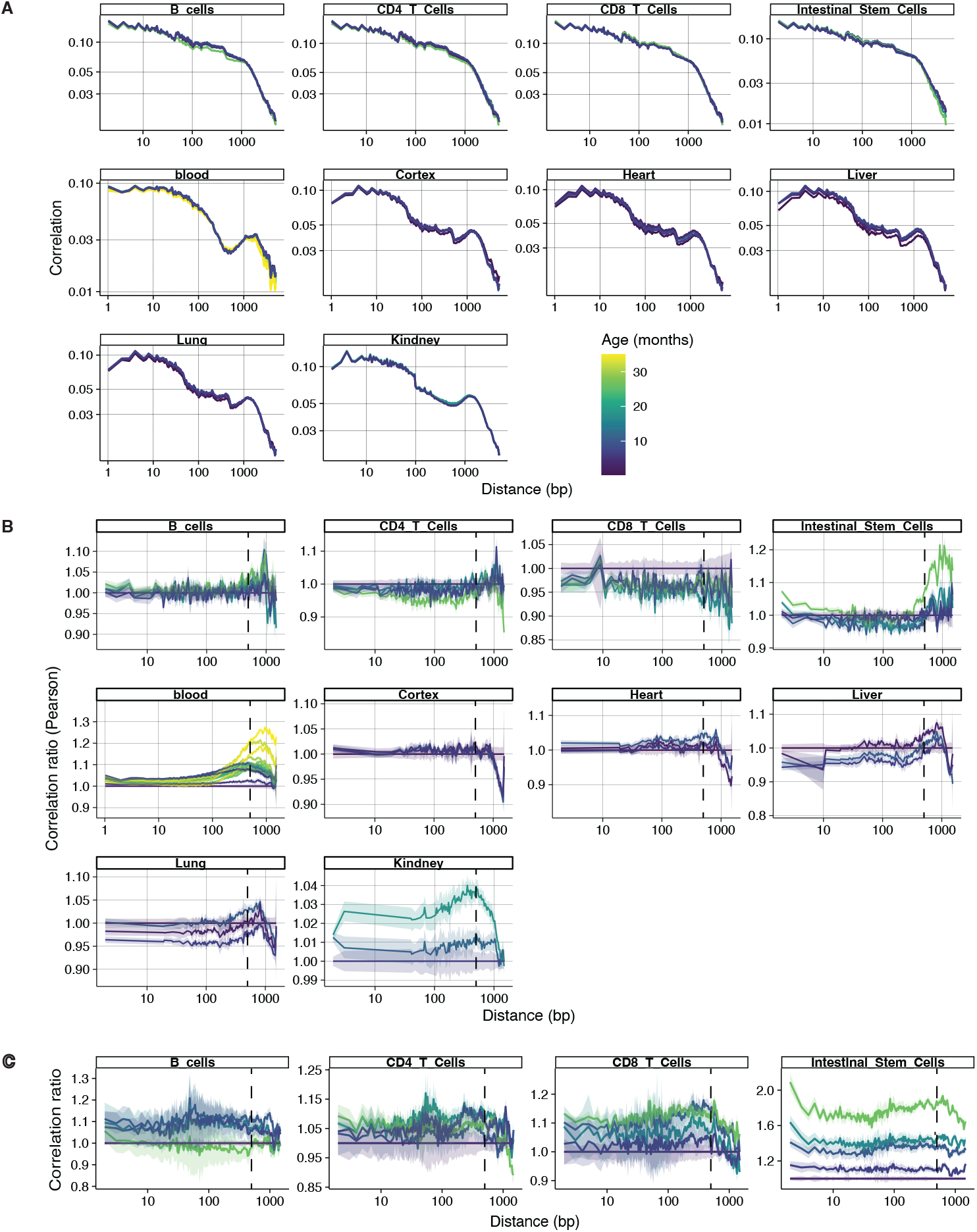
**A** Correlation functions of all analysed tissues. **B** Correlation ratio obtained from Pearson correlation for all analysed tissues. **C** Correlation ratio for tissues not shown in Figure 1.

**Figure S2:**
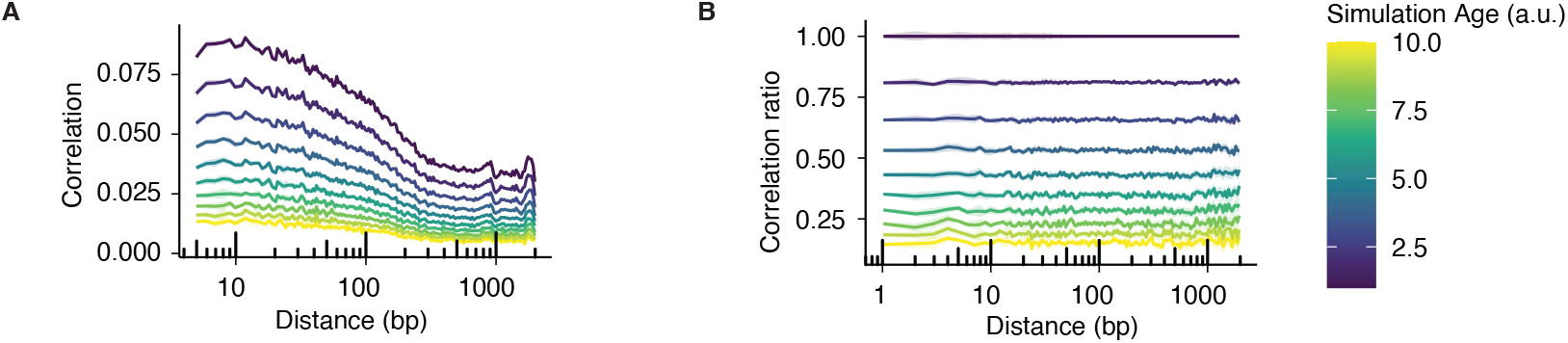
**A** Correlation for a stochastic simulation in which noise has been added to the experimental measurement of DNA methylation values in the youngest age in mouse blood [16]. **B** Correlation ratios obtained from correlation functions in **A**.

**Figure S3:**
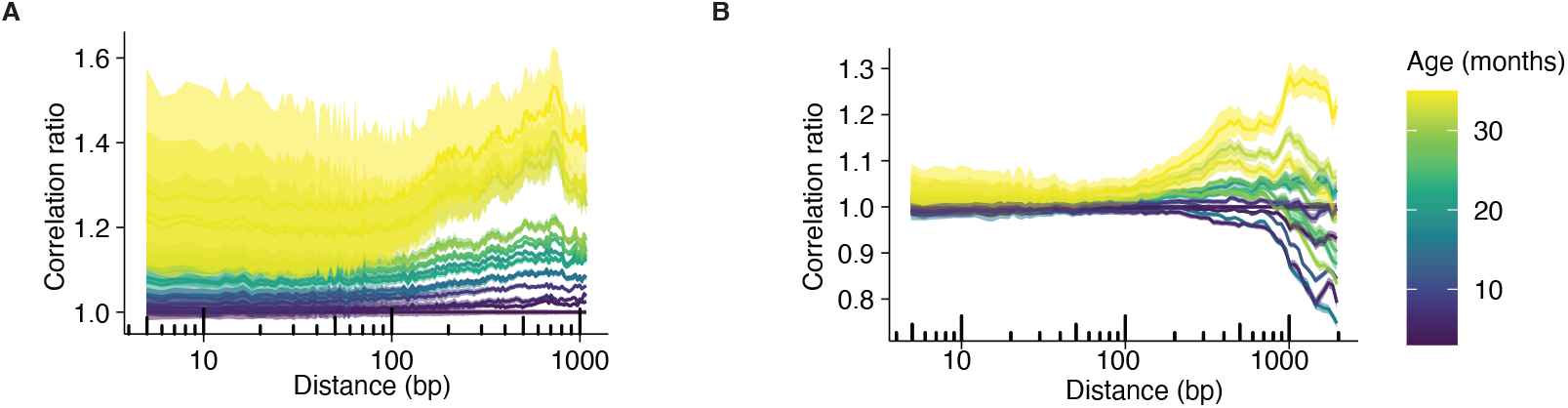
**A** Correlation ratio for CpG islands overlapping with only SUZ12 peaks. **B** Correlation ratio for CpG islands not overlapping with SUZ12 peaks.

